# Cryo-EM structures of kalium channelrhodopsins KCRs

**DOI:** 10.1101/2022.11.09.515798

**Authors:** Mingfeng Zhang, Yuanyue Shan, Li Xiao, Liping Zhao, Duanqing Pei

**Affiliations:** Laboratory of Cell Fate Control, School of Life Sciences, Westlake University, Hangzhou, 310000, China; Fudan University, Shanghai, 200433, China

## Abstract

*H. catenoides* kalium channelrhodopsins (*Hc*KCR1 and *Hc*KCR2) are potassium-selective light-gated ion channels with unique properties for optogenetics. Here we report cryo-electron microscopy (cryo-EM) structures of *Hc*KCR1 and *Hc*KCR2 with endogenous retinal. Strikingly, one structure of HcKCR1 represents the conducting state with continuous potassium ion flow in the lumen, while another in non-conducting state with discontinuous potassium ion flow. Further electrophysiological studies identified extracellular side of ion permeation region as the major element for channel potassium selectivity and retinal binding region for channel gating. Our results uncover the unprecedented potassium selectivity and light-gating mechanism, and may facilitate the rational design of new optogenetic tools.

## Introduction

Channelrhodopsins have emerged as powerful optogenetic tools to manipulate cellular ion control^1^. For example, ChR2^2^ from Chlamydomonas reinhardtii and GtACR1^3^ from Guillardia theta have been widely used for cellular cation and anion control to mediate neuron excitability or silence. Protein engineering based on the structural information of channelrhodopsins have improved their application in optogenetics. The recently identified *H. catenoides* kalium channelrhodopsins (*Hc*KCR1 and *Hc*KCR2) are of interest due to their high selectivity for K+ and unique gating by light with unprecedented fast kinetics^4^. *Hc*KCR1 in particular can deliver powerful inhibition to excitable cell firing in mouse cortical neurson by light^4^. However, these KCRs lack the canonical conserved sequence “T(S)VGY(F)G”, known as K+ selectivity filter^5^. To date, the structural basis of K+ selectivity and the gating mechanism of KCRs is unknown. Here we report the high resolution cryo-EM structures of *Hc*KCR1 and *Hc*KCR2, illuminating mechanisms for both K+ selectivity and light gating.

## Results

### *Hc*KCR1 and *Hc*KCR2 structures through CryoEM

To gain insight into these exciting KCRs, we overexpressed C-terminal GFP tagged *Hc*KCR1 and *Hc*KCR2 in HEK293T cells and performed the whole cell patch clamp under the physiological pH. The 530 nm light source was generated by the commercial Nikon fluoresce microscopy. The *Hc*KCR1 or *Hc*KCR2 transfected cells can generate reproducible and giant potassium currents by light consistent with the previous report^4^. The channel currents show a C-type inactivation^6^ like manner with the reversal potential of −70.3 ± 2.8 mV at peak photocurrent (Fig. 3d). The inactivation time from the peak current to the steady current is 33 ± 3.2 ms (Fig. 4e), and the closure time is 40.7 ± 2.2 ms (Fig. 4f). These results encouraged us to purify both *Hc*KCR1 and *Hc*KCR2 in digitonin environment (Extended Data Fig. 1a-d) and then subjected them into the routine cryo-electron microscopy (cryo-EM) process. After sample preparation, image acquisition and data processing, we obtained two near atomic *Hc*KCR1 maps of 3.17 Å and 3.35 Å resolution respectively (Extended Data Fig. 2). Meanwhile, we got 3.15 Å and 3.33 Å resolution maps of *Hc*KCR2 (Extended Data Fig. 3). The high-resolution maps of *Hc*KCR1 and *Hc*KCR2 allowed us to build molecular models of amino acids as well as the lipids and ions/waters (Extended Data Fig. 4-7).

### Trimeric Architectures of KCRs

Overall, KCRs adopt trimer architectures similar to ChRmines^7,8^, instead of dimer formation^9,10^ known for light activated non-selectivity cation channels (Fig. 1a-f). Each monomer consists of the canonical rhodopsin topology with seven transmembrane helices (TM1-TM7), three intracellular linkers (ICL1-ICL3), three extracellular linkers (ECL1-ECL3), an extracellular N-terminal domain and an intracellular C-terminal domain (Fig. 1g-h). Surprisingly, unlike ChRmines that utilize one lipid layer in the intracellular side to form the central pseudo-pore, KCRs possess two-layer lipids (Fig. 2a-c). Each monomer is connected by amino acid interaction in the trimer interface and lipids in the central cavity (Fig. 2a-c). In the extracellular side of pseudo-pore, the N-terminal loop inserts into the pseudo-pore, stabilizing trimer formation (Fig. 2d-e). In addition, the lipids of extracellular layer insert into the trimer interface formed by TM2 and TM3 and the TM4 of adjacent subunit (Fig. 2f-g). Our purified protein contains the endogenous all-trans retinal (Fig. 1a-f), which is consistent with the electrophysiological experiments that *Hc*KCRs can work without additional all-trans retinal supplementation. Therefore, our structures may present the natural functional states of *Hc*KCRs

**Fig. 1.**
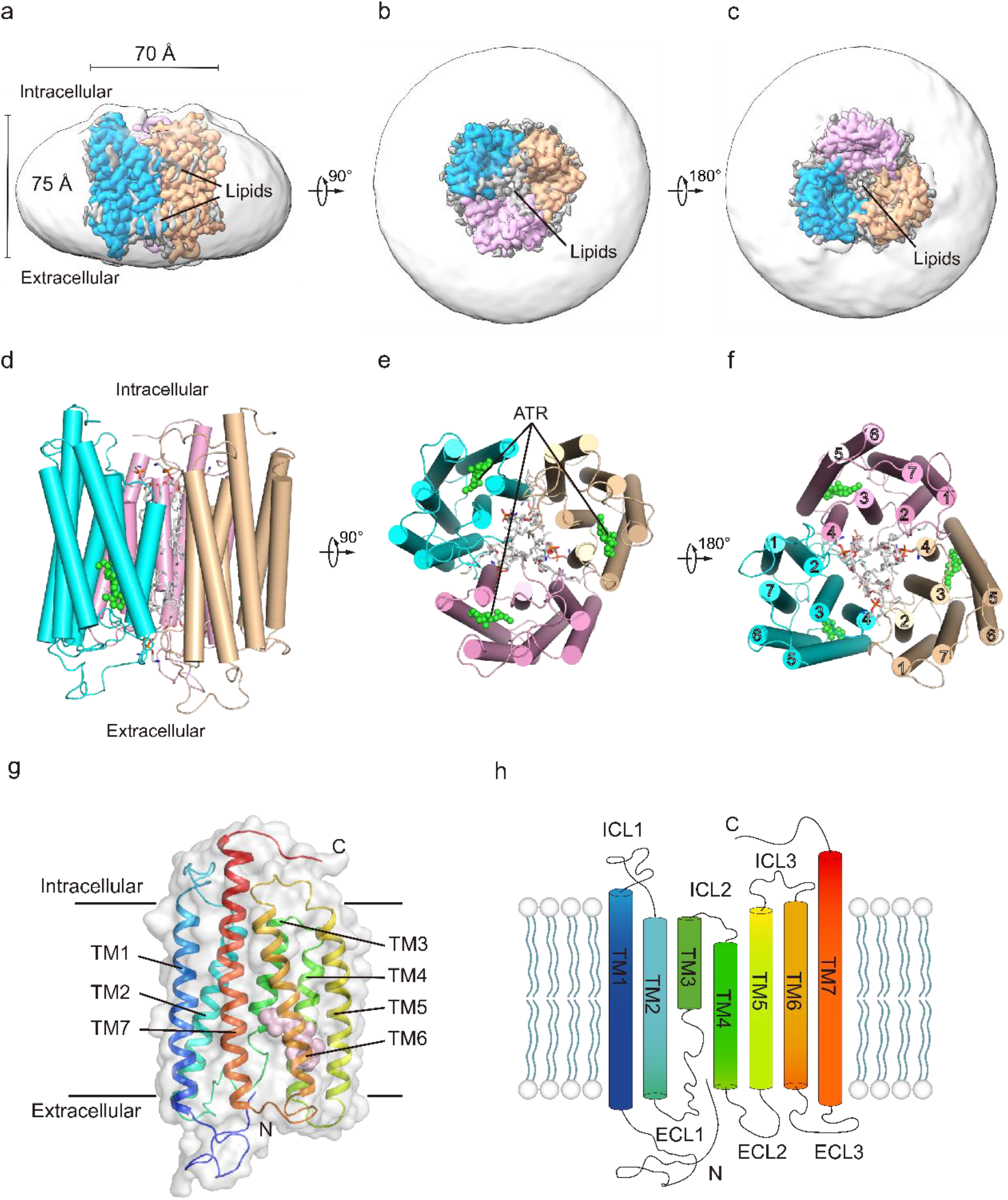
Overall structure of trimeric *Hc*KCR1. **a-c**, Cryo-EM map of trimeric *Hc*KCR1 bounded with endogenous retinal at 3.17 Å resolution viewed from the membrane plane (**a**), the extracellular side (**b**), and the intracellular side (**c**). Subunits are colored teal, pink, or wheat; the central pore lipids are colored grey; and the detergent micelle is transparent. **d-f**, Model shown in the same views as a-c respectively with endogenous all-trans retinal (ATR) shown as green spheres. **g-h**, Model (**g**) and cartoon (**h**) of a single *Hc*KCR1 subunit colored blue (N-teriminus) to red (C-terminus).

**Fig. 2.**
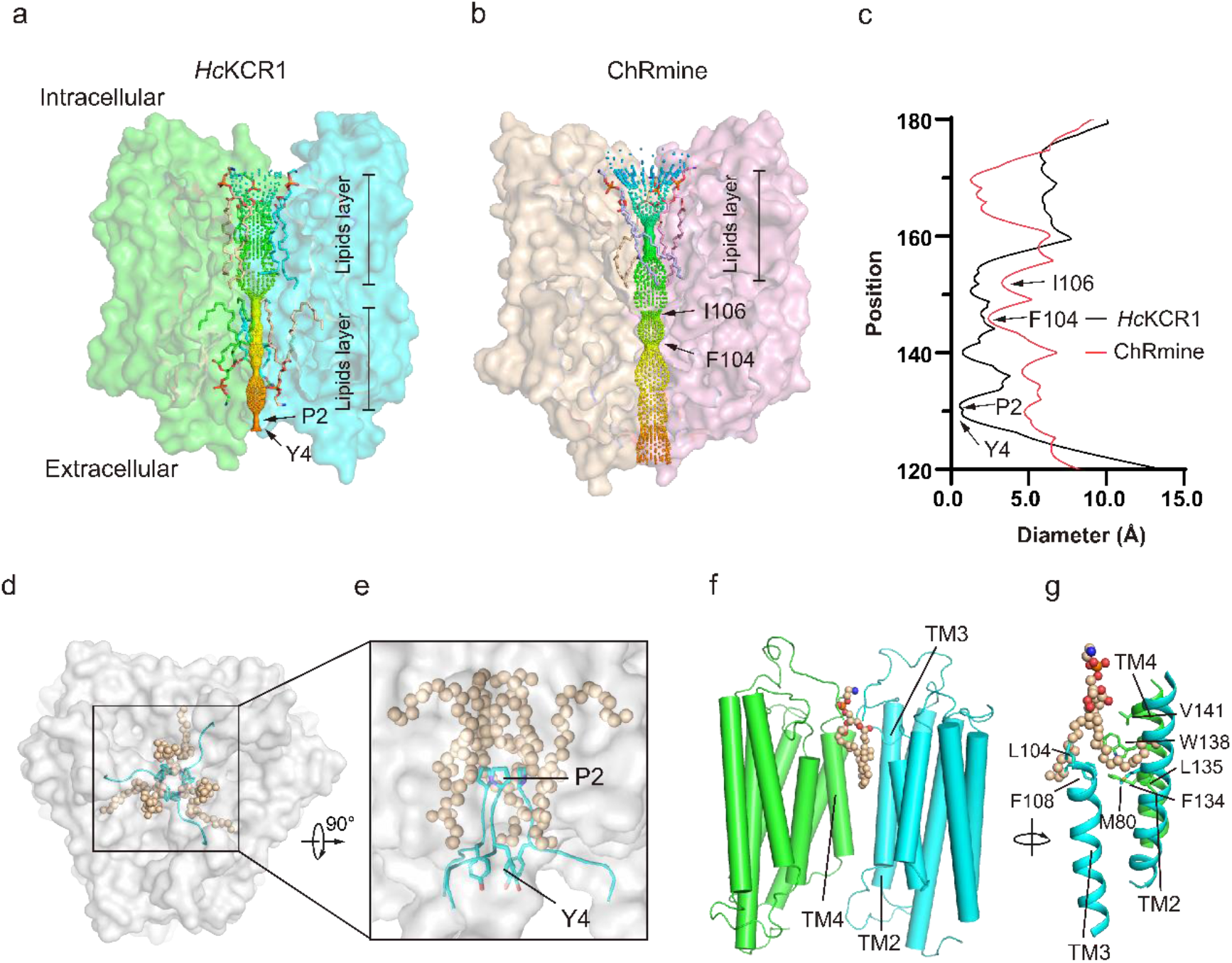
Trimeric organization of *Hc*KCR1. **a-b**, The *Hc*KCR1 pseudo-pore with two lipid layers (a) and the ChRmine pseudo-pore with one lipid layer (b) against the surface of two channel subunits (the front subunit is not displayed) viewed from the membrane plane. The restriction site, P2 and Y4 in *Hc*KCR1, I106 and F104 in ChRmine, are indicated by arrows. **c**, Pseudo-pore radius of *Hc*KCR1 and ChRmine as a function of distance from the intracellular side is shown. **d-c**, Constrictions of the pseudo-pore formed by lipids and residues (Y4 and P2) viewed form the extracellular side (**d**) and the membrane plane (**e**). **f-g**, The pseudo-pore lipids binding site in *Hc*KCR1 viewed from two angle. The key amino acids residues are indicated by arrows.

### The conducting and non-conducting states of *Hc*KCR1

The two models of overall amino acids arrangements of *Hc*KCR1 or *Hc*KCR2 are almost identical while the ion occupancies are different and retinal molecules flexible (Extended Data Fig. 6-7). We reasoned that the full ion (most-likely the potassium ions) occupancy in the lumen of *Hc*KCR1 is the conducting state and the less ion occupancy in the lumen is the non-conducting/inactivation state (Fig. 3a-b). The two maps of *Hc*KCR2 are ion-less forms (Only one or two ions can be well identified in the lumen). Since the KCRs can undergo C-type inactivation, the absence of potassium in the lumen is reminiscence of the selectivity filter for the C-type inactivation potassium channels. The previously proposed model suggests that light acts on covalently ligated all-trans retinal, leading to isomerization of retinal^11^. However, we show that, in the conducting state of HcKCR1, the retinal and K233 on TM7 do not form a canonical covalently protonated Schiff base. Instead, the density of the retinal located region seems to stand for a potassium ion locating between the K233 and the retinal moiety (Extended Data Fig. 6). Consequently, all potassium ions in the long narrow lumen form a continuous ion flow (Fig. 3a).

**Fig. 3.**
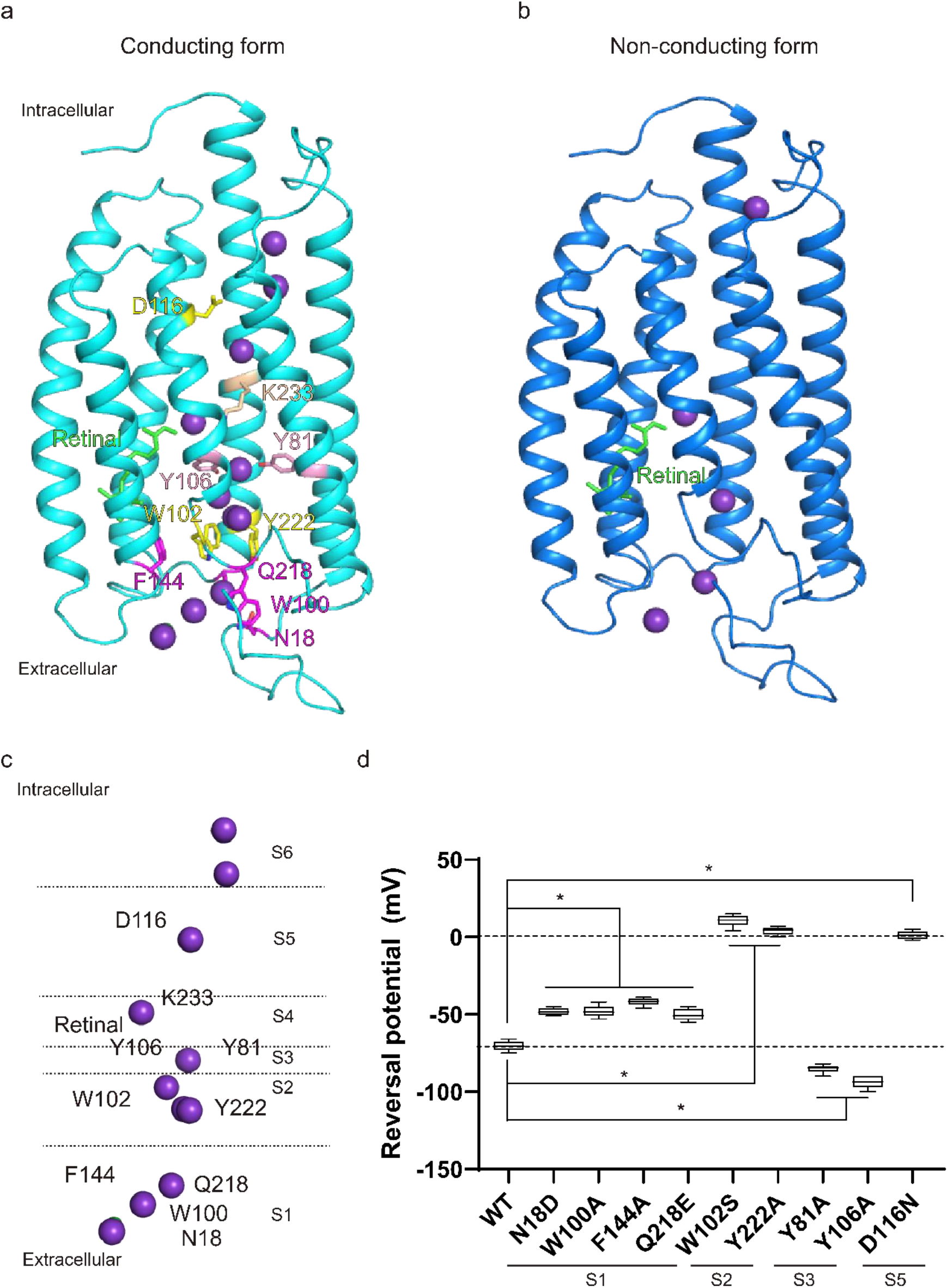
Ion selectivity of *Hc*KCR1. **a-b**, Conducting form (**a**) and non-conducting forms of *Hc*KCR1 subunit (**b**) viewed from the membrane plane. The endogenous retinal is shown as green stick and potassium ions in pore domain are shown as purple spheres. The residues interacting with potassium ion is shown in **a. c**, Ion flow in *Hc*KCR1 pore domain. The pore domain are divided into six layers (S1-S6). **d**, The reversal potentials of the peak current of wild-type *Hc*KCR1 and its mutants. The data points are the mean ± sem (n is at least 3 cells for each variant). *, p < 0.05 by the two-tailed paired sample Wilcoxon signed rank test.

### The mechanism for potassium selectivity in KCRs

We divided the continuous potassium ions occupancy region into six different layer S1-S6 as shown in Figure 3a and 3c. The mutations of residues in S1 layer resulted in slightly reduced potassium selectivity, while the mutations of residues in S2 layer led to very largely reduced potassium selectivity. Additionally, the mutations of residues in S3 layer increase the potassium selectivity. Surprisingly, the D116N mutation in S5 layer also exhibit the non-selectivity manner (Fig. 3c-d). We reasoned that the S1 and S2 layers, especially S2 layer, play vital roles in potassium selectivity with multiple π-π interactions formed by residues with phenyl, hydroxyphenyl or indole rings, such as W100, F144, W102 and Y222. Furthermore, the potassium and the phenyl, hydroxyphenyl or indole rings may form a π-cation interaction for specific potassium selectivity. In the S3 layer, two tyrosine residues form a gate-like barrier. Abolishing the side chain of this two tyrosine promotes the potassium fast flow consequently increases the potassium ion selectivity. Unexpectedly, electrophysical results of D116N in S5 layer exhibit the non-selectivity manner, which indicates that the D116 may also function as the potassium filter or have indirectly allosteric influence on S1 and S2 regions.

### The gating kinetics of KCRs

To explore the potential mechanism of gating kinetics induced by light, we screened the mutations in retinal binding site and the S4 layer region (Fig. 4a-c). The mutation of retinal-coordinated residue W119L largely increase the inactivation rate while no significant changes in the closure rate and potassium selectivity (Fig. 4d-g). The mutation of S4 layer coordinated residue T109V largely attenuates the closure and inactivation rate, while no significant changes in potassium selectivity (Fig. 4d-g). These results suggest that retinal receives the light signal that transit to W119 and T109 for channel gating kinetics.

**Fig. 4.**
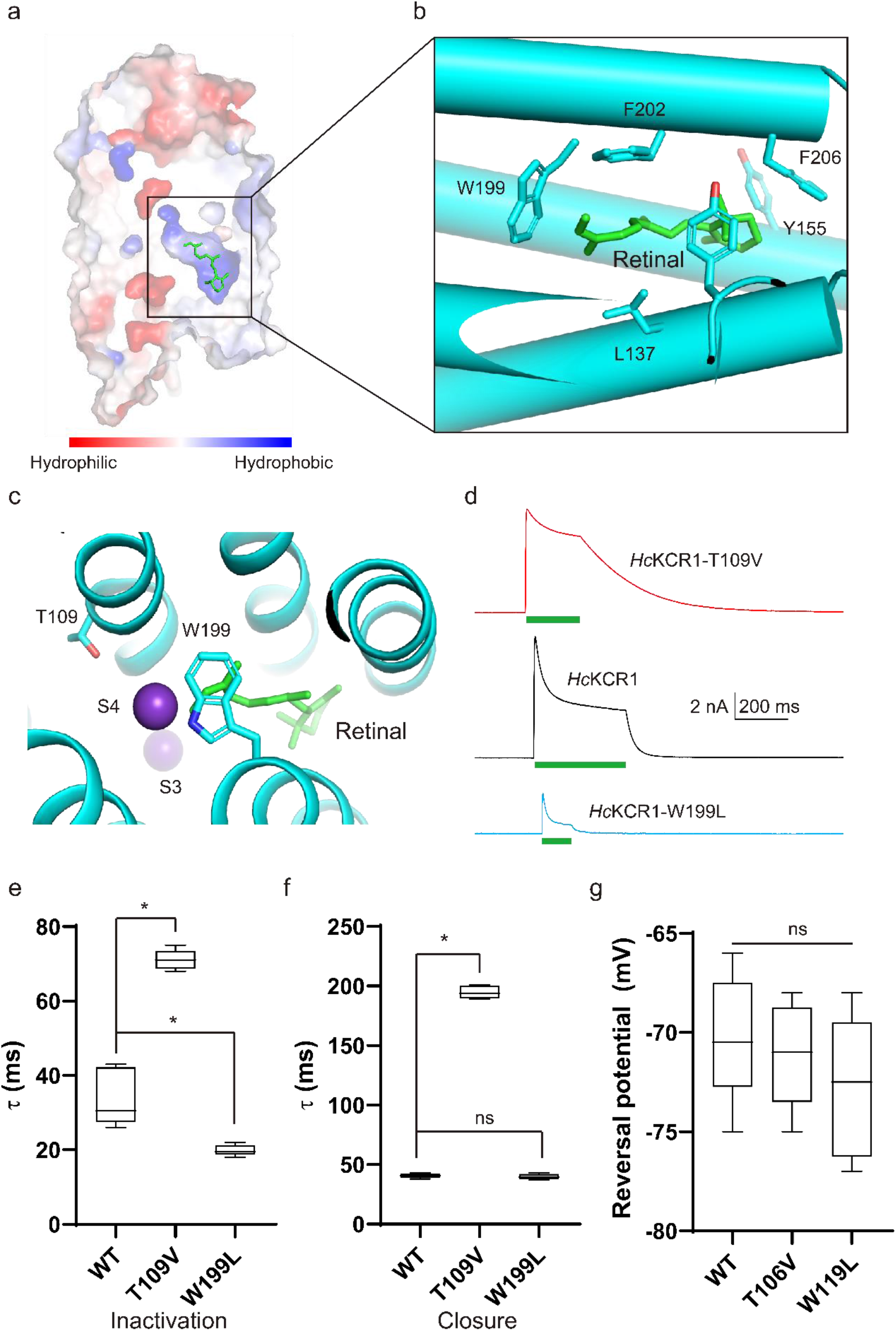
Light-gating kinetics of *Hc*KCR1. **a**, Electrostatic potential surface of *Hc*KCR1. **b**, Enlarged view of all-trans retinal binding site in *Hc*KCR1. The related transmembrane helices are shown as cartoon and the side chain of key residues are shown as sticks. **c**, Enlarged view of *Hc*KCR1 light-gating site. Potassium ion are shown as purple spheres and retinal is shown as green sticks. **d**, Representative photocurrents at 0 mV voltage for *Hc*KCR1 and each mutant evaluated. The green bar under each trace indicates the light pulse. **e-f**, Inactivation decay time (**e**) and closed decay time (**f**) of *Hc*KCR1, T109V, and W199L. **g**, The reversal potentials of the peak current of wild-type *Hc*KCR1, T109V, and W199L. The data points are the mean±sem (n is at least 3 cells for each variant). *, p < 0.05 by the two-tailed paired sample Wilcoxon signed rank test.

## Discussion

In *Hc*KCRs, the potassium permeation pathway is a long and narrow hydrophilic lumen across the whole cell membrane and gated by the covalently protonated Schiff base formed by the all trans retinal chromophore and the conserved lysin. Our model of the conducting state suggested that the light can break up the covalently protonated Schiff base and then open the channel.

Unlike the golden principle of the channel potassium selectivity with conserved sequence “T(S)VGY(F)G”, our results indicate that multiple residues in the permeation path are involved in channel potassium selectivity. The pivotal residue is the W102, which is likely to act as a central role connecting multiple π-π interactions between the phenyl, hydroxyphenyl or indole rings containing residues and π-cation interactions between potassium and the phenyl, hydroxyphenyl or indole rings containing residues. This new principle of potassium selectivity may provide rationales to design organic artificial potassium channels^12^.

The mutations around gate region may allow us to discover better optogenetic tools, including the high potassium selectivity mutations (Y81A and Y106A). In addition, the W119L has a desirable property with a fast inactivation kinetics. Further more useful optogenetic tools may be engineered based on further high resolution and time resolved conformations of *Hc*KCRs.

In sum, we solved the structures of *Hc*KCRs which illuminate key mechanisms for potassium-selectivity and light-gating unique to this family of channels. These results may further encourage the development of these ChRs into tools for optogenetics.

## Methods

### Cloning and protein expression

The coding sequence for KCR1 and KCR2 from *Hyphochytrium catenoides* were cloned into a custom vector based on the BacMam expression backbone with an added C-termainl PreScission protease (PPX) cleavage site, linker sequence, enhanced GFP (EGFP), and a FLAG tag to generate the constructs KCR1/2-LE-LEVLFQGP-GGK-EGFP-GSG-DYKDDDDK for expression. Mutations were introduced using inverse PCR.

The *Hc*KCR1/2 constructs were transformed into DH10Bac *E*.*coli* to generate a bacmid according to the manufacturer’s instructions. Subsequent steps used *Spodopetera frugiperda* sf9 cells cultured in sf-900 II SFM medium (Thermo Fisher Scientific). Bacmid was transfected into adherent sf9 cells using the cellfectin (Gibco) to produce P1 virus. P1 virus was used to infect sf9 cells in suspension at 1.5 million cells/mL at a multiplicity of infection (MOI) ∼0.1 to generate P2 virus. Infection was monitored by fluorescence and P2 virus was harvested 48-72 hours post infection. P3 virus was generated in a similar manner. P3 viral stock was then used to infect expi-HEK293F cells at 2.5 million cells/mL at a MOI ∼2-5 for large scale protein expression. Cells were harvested by centrifugation at 4000×g for 15 min, flash-frozen in liquid nitrogen, and stored at −80°C.

### Protein purification

Cells from 1 L of culture were thawed and resuspended in 100 mL of lysis buffer (20 mM Tris-HCl, 150 mM KCl, pH 8.0). Cells were lysed by sonication and membranes pelleted by centrifugation at 40,000 r.p.m. for 40 min in a Ti45 rotor. Supernatant was load onto anti-FLAG resin (Genscript) by gravity flow. Resin was further washed with 10 column volumes of wash buffer (lysis buffer plus 0.02% LMNG), and protein was eluted with an elution buffer (wash buffer plus 230 μg/ml FLAG peptide). The C-terminal GFP tag of eluted protein was removed by HRV3C protease cleavage for 2 h at 4°C. The protein was further concentrated by a 100-kDa cutoff concentrator (Milipore) and loaded onto a Superose 6 increase 10/300 column (GE Healthcare) running in lysis buffer plus 0.03% digitonin. Peak fractions were combined and concentrated to around 9 mg/ml of *Hc*KCR1 and 8 mg/ml of *Hc*KCR2 for cryo-EM sample preparation.

### EM sample preparation

The protein samples were cleared by centrifugation at 35,000 r.p.m. for 30 min at 4°C prior to grid preparation. A 3-μl drop of protein was applied to freshly glow discharged Holey Carbon, 300 mesh R1.2/1.3 gold grids (Quantifoil). Sample were plunge frozen in liquid nitrogen cooled liquid ethane using a FEI Vitrobot Mark IV (Thermo Fisher Scientific) at 8°C, 100% humidity, 3 blot force, ∼5 s wait time, and 6 s blot time.

### Cryo-EM data acquisition

Cryo-EM data for *Hc*KCR1/2 were collected on a Titan Krios microscope (FEI) equipped with a cesium corrector operated at 300 kV. Movie stacks were automatically acquired with EPU software on a Gatan K3 Summit detector in super-resolution mode (105,000× magnification) with pixel size 0.4235 Å at the object plane and with defocus ranging from −1.4 μm to −1.9 μm and GIF Quantum energy filter with a 20 eV slit width. The does rate on the sample was 23.438 e^-^ s^-1^ Å^-2^, and each stack was 2.56 s long and does-fractioned into 32 frames with 80 ms for each frame. Total exposure was 60 e^-^ Å^-2^. Acquisition parameters are summarized in table S1.

### Cryo-EM data processing

A flowchart of the Cryo-EM data processing process can be found in Fig. S2. Data processing was carried out with the cryoSPARC v3 suite. Super-resolution image stacks were gain-normalized, binned by 2 with Fourier cropping, and patch-based CTF parameters of the dose-weighted micrographs (0.849 Å per pixel) were determined by cryoSPARC.

For dataset of HcKCR1, around 607K particles were picked by blob picking from 999 micrographs and 2D classification was performed to select 2D classes as templates. Template picker was used to pick 1,016K particles. Particles were extracted using a 320-pixel box with a pixel size of 0.849 Å. The dataset was cleared using several rounds of 2D classification, followed by Ab initio reconstruction using C1 symmetry and initial model (PDB: 7SFK). Two maps with 94.7K (22.3%) and 102.6K (24.5%) particles were refined using non-uniform refinement and local refinement for reconstructing the density map while imposing a C3 symmetry. The two final maps were low-pass filtered to the map-model FSC value. The reported resolutions were based on the FSC=0.143 criterion.

For dataset of HcKCR2, around 598K particles were picked by blob picking from 999 micrographs and 2D classification was performed to select 2D classes as templates. Template picker was used to pick 1,144K particles. Particles were extracted using a 320-pixel box with a pixel size of 0.849 Å. The dataset was cleared using several rounds of 2D classification, followed by Ab initio reconstruction using C1 symmetry and initial model (PDB: 7SFK). Two maps with 171.3K (21.9%) and 173.8K (23.2%) particles were refined using non-uniform refinement and local refinement for reconstructing the density map while imposing a C3 symmetry. The two final maps were low-pass filtered to the map-model FSC value. The reported resolutions were based on the FSC=0.143 criterion.

### Model building

The atomic models of monomer were built in Coot based on an initial model (PDB: 7SFK). The models were then manually adjusted in Coot. Trimeric models were obtained subsequently by applying a symmetry operation on the monomer. These trimeric models were refined using phenix.real_space_refine with secondary structure restraints and Coot iteratively. Residues whose side chains have poor density were modeled as alanines. For validation, FSC curves were calculated between the final models and EM maps. The pore radii were calculated using HOLE. Figures were prepared using PyMOL and Chimera.

### Whole-cell patch clamp recording from HEK293 cells

The HEK293 cells from the American Type Culture Collection (ATCC) were were cultured on coverslips placed in a 12-well plate containing DMEM/F12 medium (Gibco) supplemented with 10% fetal bovine serum (FBS). The cells in each well were transiently transfected with 1 μg *Hc*KCR1/2-CGFP DNA plasmid using 3μg linear polyethylenimine (PEI, MW 25000, Polysciences) according to the manufacturer’s instructions. After 12-20 h, the coverslips were transferred to a recording chamber containing the external solution (10 mM HEPES-Na pH 7.4, 150 mM NaCl, 5 mM glucose, 2 mM MgCl_2_ and 1 mM CaCl_2_). Borosilicate micropipettes (OD 1.5 mm, ID 0.86 mm, Sutter) were pulled and fire polished to 4-6 MΩ resistance. The pipette solution was 10 mM HEPES-Na pH 7.4, 150 mM KCl, 5 mM EGTA. The bath solution was 10 mM HEPES-Na pH 7.4, 150 mM NaCl, 5 mM glucose, 2 mM MgCl_2_ and 1 mM CaCl_2_. All measurements were carried out at room temperature (∼25°C) in dark environment using an Axopatch 200B amplifier, a Digidata 1550 digitizer and pCLAMP software (Molecular Devices). A light source built in commercial Nikon fluoresce microscopy with a mechanical shutter was used as the light source (maximal quantum density at the focal plane of the 40× objective lens was ∼7 mW mm^-2^ at 530 nm). The patches were held at −60 mV and the recordings were low-pass filtered at 1 kHz and sampled at 20 kHz. All current-voltage curve (IV dependencies) were corrected for liquid junction potentials (LJP) calculated using the ClampEx built-in LJP calculator. Statistical analyses were performed using ClampFit and Graphpad Prism.

## Data availability

The coordinate files and cryo-EM maps of the conducting *Hc*KCR1, non-conducting *Hc*KCR1, class I *Hc*KCR2 and class II *Hc*KCR2 have been deposited at Protein Data Bank (Electron Microscopy Data Bank (EMDB)) under accession codes 8H22 (EMD-34435), 8H23 (EMD-34436), 8H1X (EMD-34433) and 8H1Z (EMD-34434), respectively.

## Acknowledgements

We would like to thank the Cryo-EM Facility of Westlake University for providing cryo-EM and computation support. This work was supported by Westlake Laboratory (Westlake Laboratory of Life Sciences and Biomedicine) and an Institutional Startup Grant from the Westlake Education Foundation to D.P. We also would like to thank all the Cell fate control lab members for their support.

## Author contributions

M.Z. and D.P conceived the project. Y.S. and M.Z. designed the experiments. Y.S. prepared the cryo-EM sample and performed electrophysiology experiments. Y.S. and M.Z. collected cryo-EM data. M.Z. performed image processing and analyzed EM data. Y.S. and M.Z. built the model and wrote the manuscript draft. All authors contributed to the manuscript preparation.

## Conflict of interest

The authors declare no conflict of interest.

**Extended Data Fig.1.**
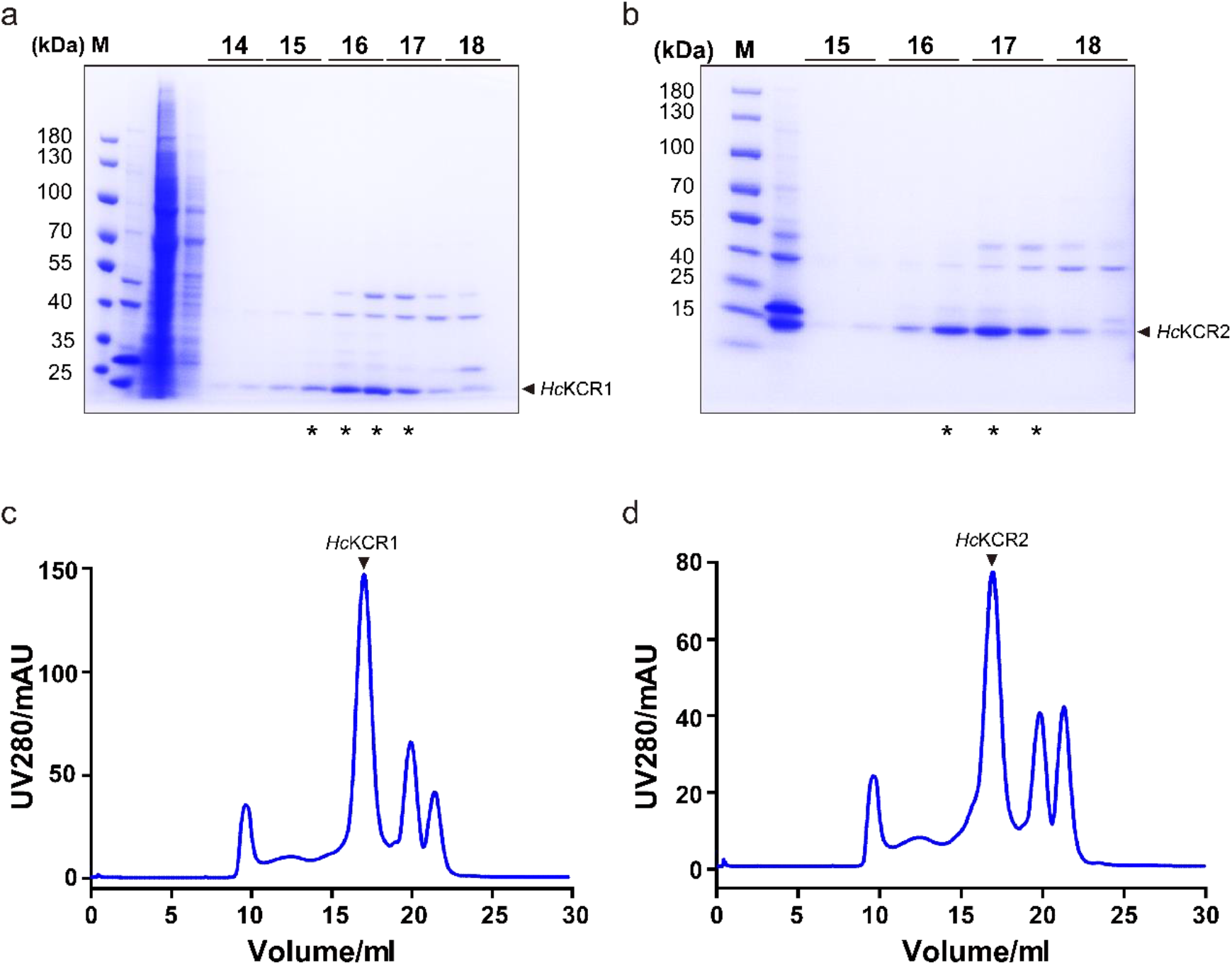
Purification of the *Hc*KCR1 and *Hc*KCR2. **a-b**, SDS-PAGE gel of peak fractions of *Hc*KCR1 (**a**) and *Hc*KCR2 (**b**) stained with Coomassie blue. The predicted molecular weight based on protein sequence for a single KCR1/2 subunit is around 29.4 kDa. **c-d**, Representative size-exclusion chromatography (SEC) trace of purified *Hc*KCR1 (**c**) and *Hc*KCR2 (**d**).

**Extended Data Fig.2.**
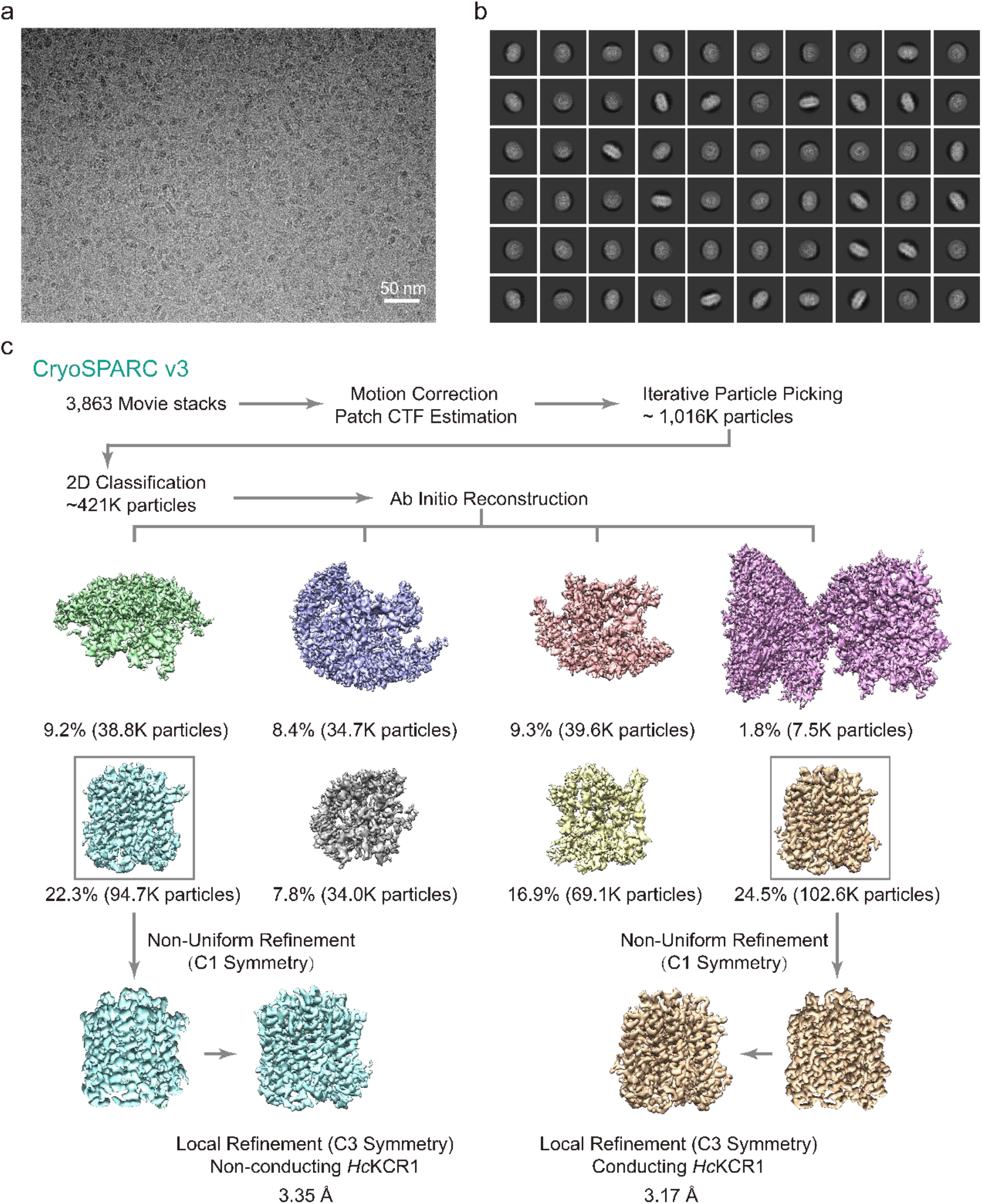
Single-particle Cryo-EM reconstructions of the *Hc*KCR1. **a**, A representative raw micrograph of the *Hc*KCR1. **b**, Selected 2D class averages. **c**, Summary of image processing for six states of *Hc*KCR1 dataset with C3 symmetry.

**Extended Data Fig.3.**
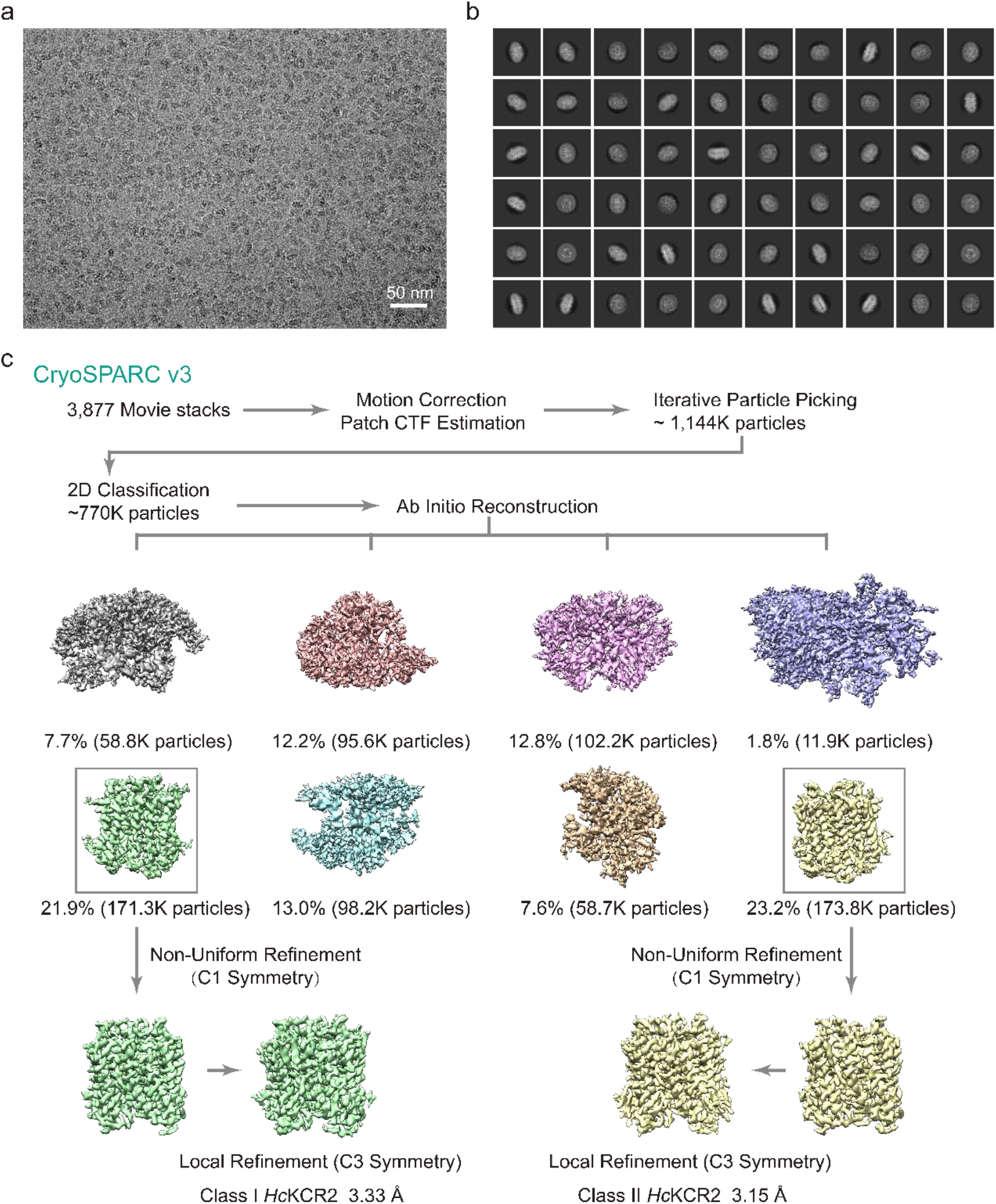
Single-particle Cryo-EM reconstructions of the *Hc*KCR2. **a**, A representative raw micrograph of the *Hc*KCR2. **b**, Selected 2D class averages. **c**, Summary of image processing for six states of *Hc*KCR2 dataset with C3 symmetry.

**Extended Data Fig.4.**
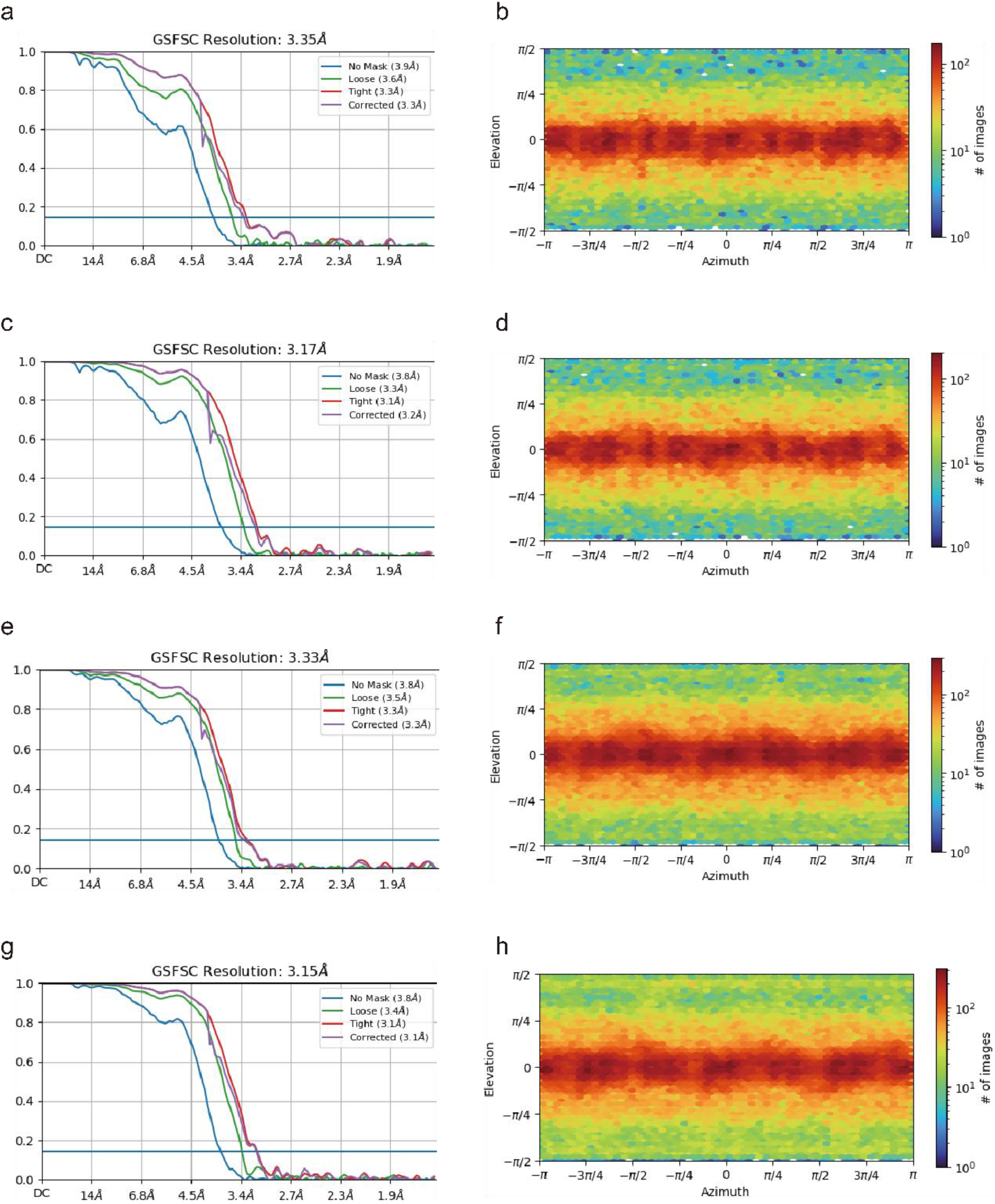
Fourier Shell correlation (FSC) curves and Euler angle distribution of particles for the final 3D reconstruction of *Hc*KCR1 and *Hc*KCR2. **a-h**, FSC curves between two half maps before (blue) and after loose (Green) and tight (Purple) masks of non-conducting *Hc*KCR1 (**a**), conducting *Hc*KCR1 (**c**), class I *Hc*KCR2 (**e**), and class II *Hc*KCR2 (**g**), respectively. Euler angle distribution of particles for 3D reconstruction of h *Hc*KCR1 in non-conducting forms (**b**), conducting form (**d**) and class I *Hc*KCR2 (**f**), class II *Hc*KCR2 (**h**), respectively. The reported resolutions were based on the FSC=0.143 criterion.

**Extended Data Fig.5.**
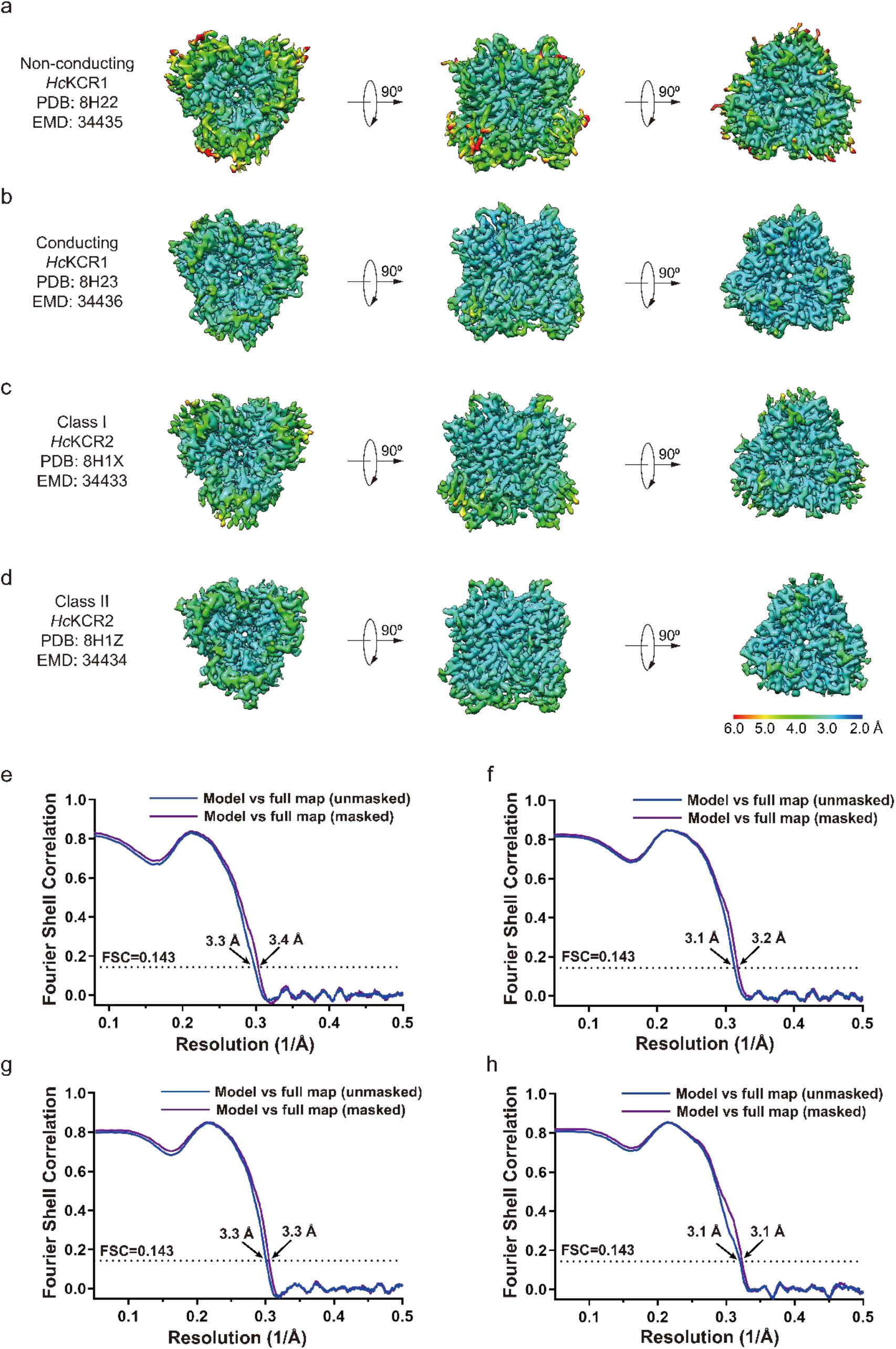
Local resolution and FSC curves for cross-validation of two density maps of *Hc*KCR1 and two density maps of *Hc*KCR2. **a-d**, Local resolution of non-conducting *Hc*KCR1 (**a**), conducting *Hc*KCR1 (**b**), class I *Hc*KCR2 (**c**), and class II *Hc*KCR2 (**d**), in bottom view (left), side view (middle) and top view (right) is shown. **e-h**. FSC curves for cross-validation: model versus unmasked full map (blue) and masked full map (purple) of non-conducting *Hc*KCR1 (**e**), conducting *Hc*KCR1 (**f**), class I *Hc*KCR2 (**g**), and class II *Hc*KCR2 (**h**).

**Extended Data Fig.6.**
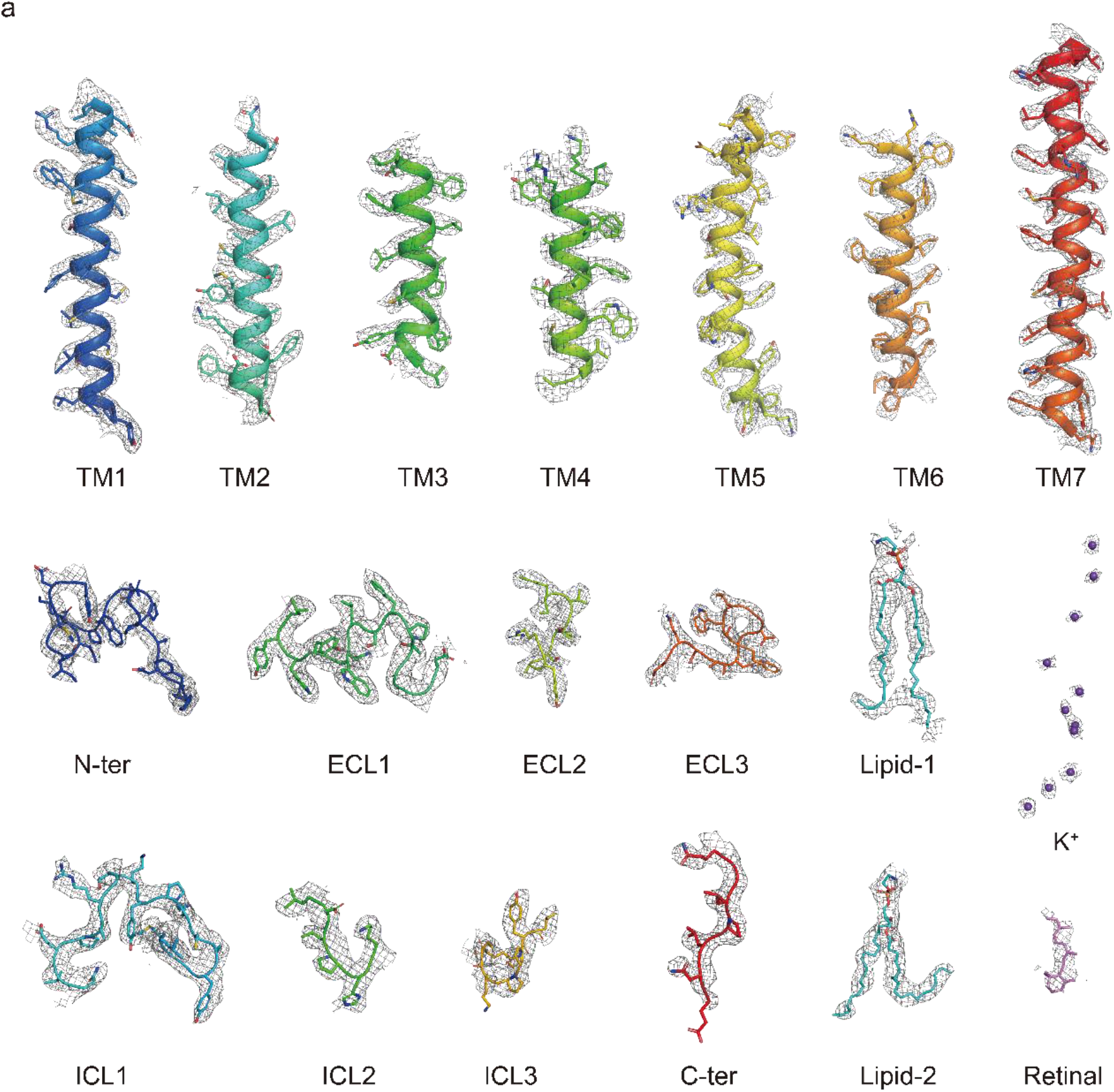
EM density and of conducting *Hc*KCR1. **a**, The cryo-EM density of *Hc*KCR1, lipids, endogenous retinal and the putative potassium ions.

**Extended Data Fig.7.**
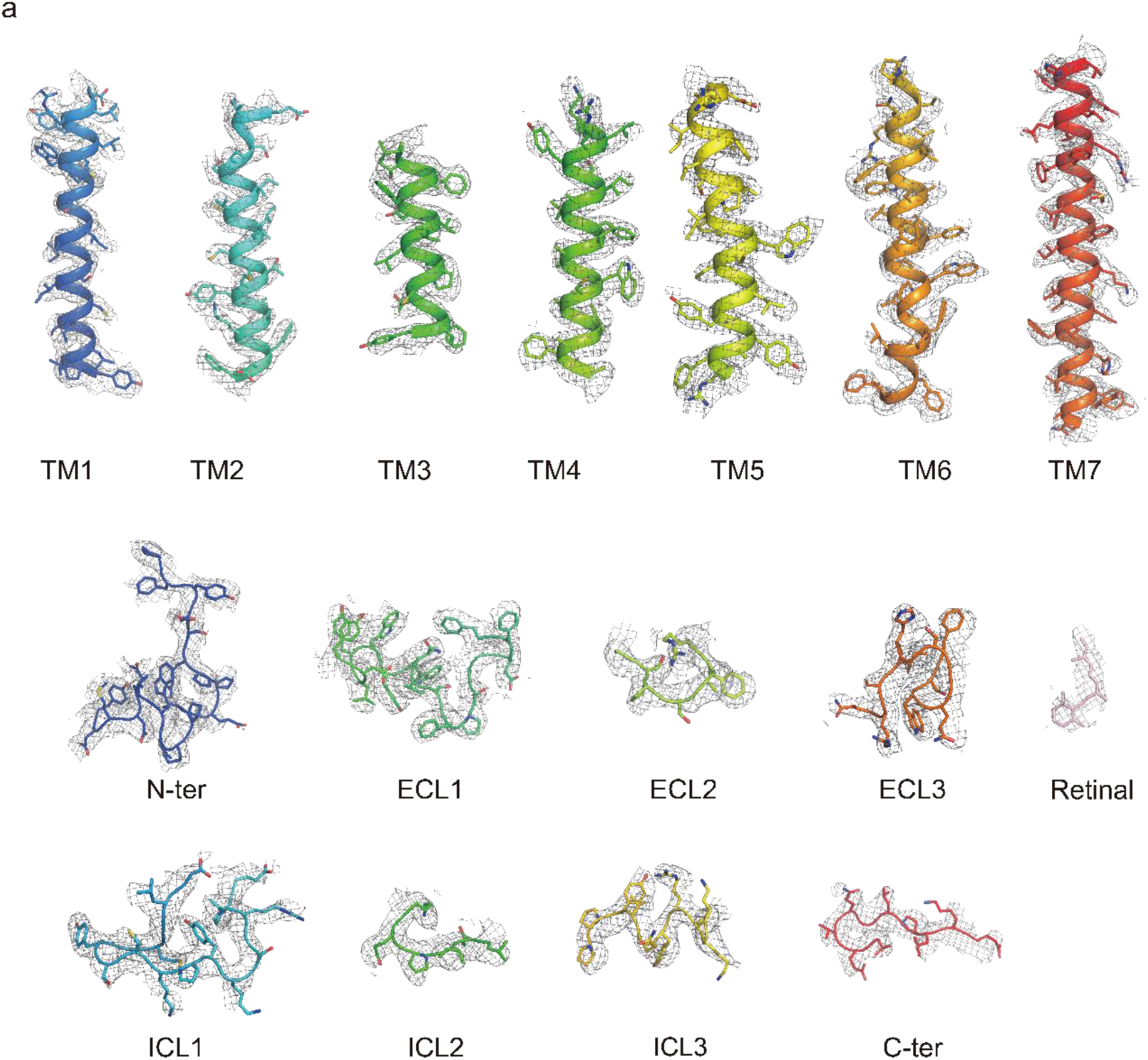
EM density and of class II *Hc*KCR2. **a**, The cryo-EM density of *Hc*KCR2 and endogenous retinal.

**Table S1.**
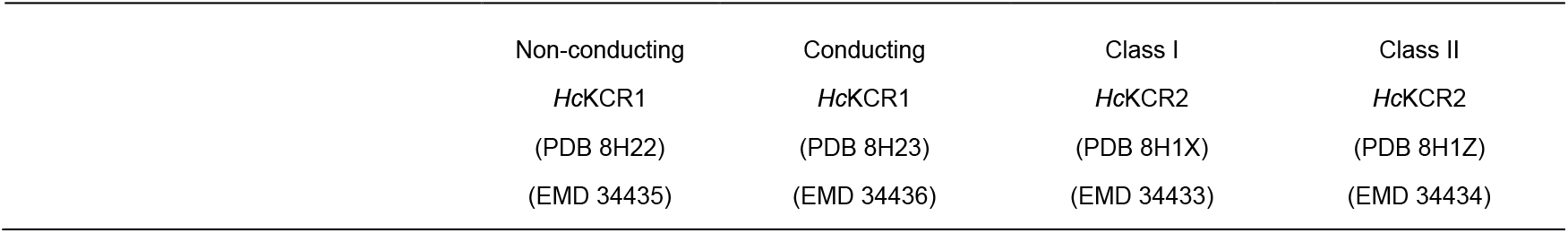
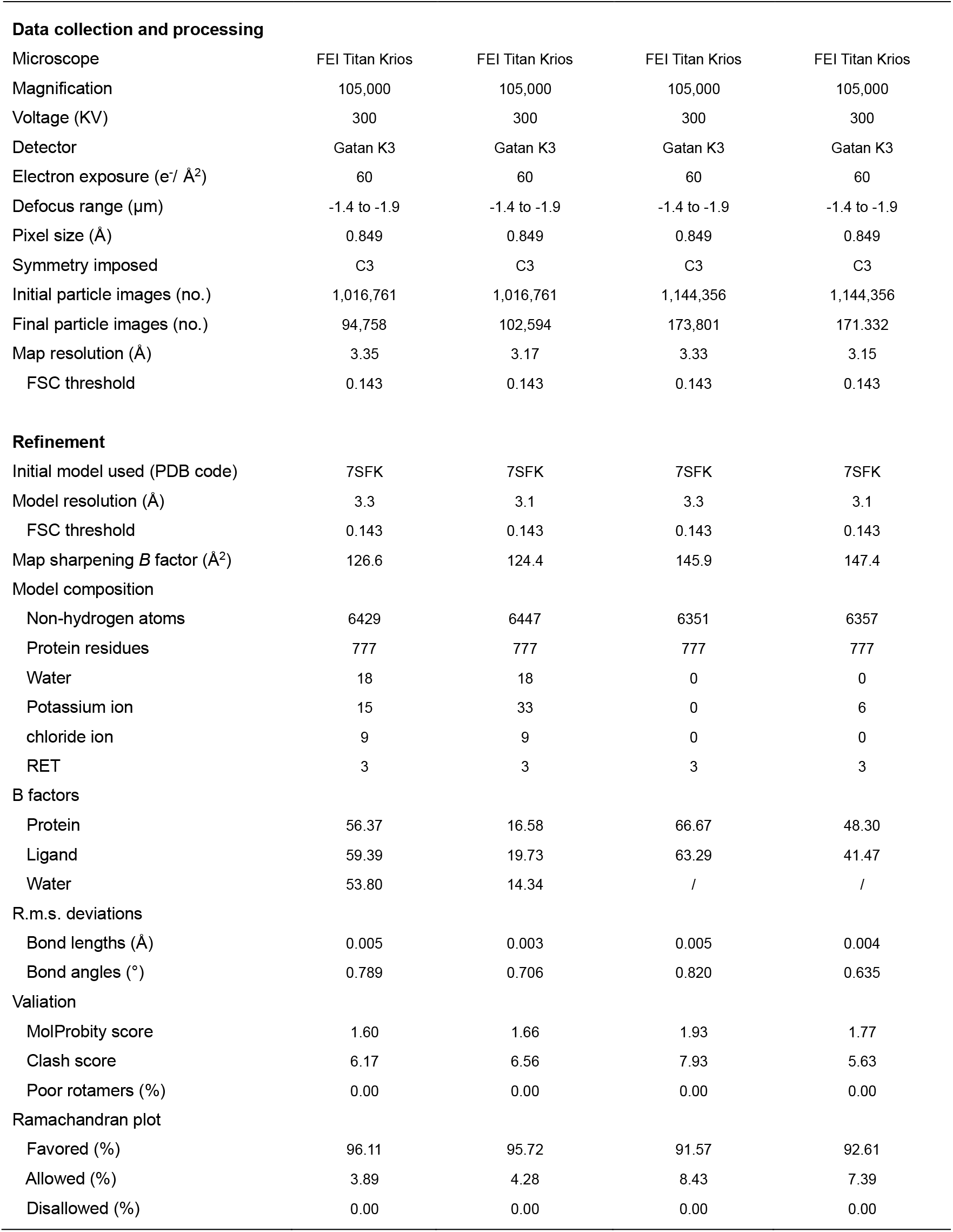
Cryo-EM data collection, refinement, and validation statistics.

| Non-conducting | Conducting | Class I | Class II |
| --- | --- | --- | --- |
| *HcKCR1* | *HcKCR1* | *HcKCR2* | *HcKCR2* |
| (PDB 8H22) | (PDB 8H23) | (PDB 8H1X) | (PDB 8H1Z) |
| (EMD 34435) | (EMD 34436) | (EMD 34433) | (EMD 34434) |

|  |  |  |  |  |
| --- | --- | --- | --- | --- |
| **Data collection and processing** |  |  |  |  |
| Microscope | FEI Titan Krios | FEI Titan Krios | FEI Titan Krios | FEI Titan Krios |
| Magnification | 105,000 | 105,000 | 105,000 | 105,000 |
| Voltage (KV) | 300 | 300 | 300 | 300 |
| Detector | Gatan K3 | Gatan K3 | Gatan K3 | Gatan K3 |
| Electron exposure (e<sup>-</sup>/ Å<sup>2</sup>) | 60 | 60 | 60 | 60 |
| Defocus range (μm) | -1.4 to -1.9 | -1.4 to -1.9 | -1.4 to -1.9 | -1.4 to -1.9 |
| Pixel size (Å) | 0.849 | 0.849 | 0.849 | 0.849 |
| Symmetry imposed | C3 | C3 | C3 | C3 |
| Initial particle images (no.) | 1,016,761 | 1,016,761 | 1,144,356 | 1,144,356 |
| Final particle images (no.) | 94,758 | 102,594 | 173,801 | 171,332 |
| Map resolution (Å) | 3.35 | 3.17 | 3.33 | 3.15 |
| FSC threshold | 0.143 | 0.143 | 0.143 | 0.143 |
| **Refinement** |  |  |  |  |
| Initial model used (PDB code) | 7SFK | 7SFK | 7SFK | 7SFK |
| Model resolution (Å) | 3.3 | 3.1 | 3.3 | 3.1 |
| FSC threshold | 0.143 | 0.143 | 0.143 | 0.143 |
| Map sharpening *B* factor (Å<sup>2</sup>) | 126.6 | 124.4 | 145.9 | 147.4 |
| Model composition |  |  |  |  |
| Non-hydrogen atoms | 6429 | 6447 | 6351 | 6357 |
| Protein residues | 777 | 777 | 777 | 777 |
| Water | 18 | 18 | 0 | 0 |
| Potassium ion | 15 | 33 | 0 | 6 |
| chloride ion | 9 | 9 | 0 | 0 |
| RET | 3 | 3 | 3 | 3 |
| B factors |  |  |  |  |
| Protein | 56.37 | 16.58 | 66.67 | 48.30 |
| Ligand | 59.39 | 19.73 | 63.29 | 41.47 |
| Water | 53.80 | 14.34 | / | / |
| R.m.s. deviations |  |  |  |  |
| Bond lengths (Å) | 0.005 | 0.003 | 0.005 | 0.004 |
| Bond angles (°) | 0.789 | 0.706 | 0.820 | 0.635 |
| Valiation |  |  |  |  |
| MolProbity score | 1.60 | 1.66 | 1.93 | 1.77 |
| Clash score | 6.17 | 6.56 | 7.93 | 5.63 |
| Poor rotamers (%) | 0.00 | 0.00 | 0.00 | 0.00 |
| Ramachandran plot |  |  |  |  |
| Favored (%) | 96.11 | 95.72 | 91.57 | 92.61 |
| Allowed (%) | 3.89 | 4.28 | 8.43 | 7.39 |
| Disallowed (%) | 0.00 | 0.00 | 0.00 | 0.00 |

